# Targeted cell ablation in zebrafish using optogenetic transcriptional control

**DOI:** 10.1101/730507

**Authors:** Karen Mruk, Paulina Ciepla, Patrick A. Piza, Mohammad A. Alnaqib, James K. Chen

**Affiliations:** Department of Chemical and Systems Biology, Stanford University School of Medicine, Stanford, CA, USA; Department of Developmental Biology, Stanford University School of Medicine, Stanford, CA, USA; School of Pharmacy, University of Wyoming, Laramie, WY, USA

**Keywords:** Optogenetics, nitroreductase, M2 ion channel, cell ablation, neural injury, zebrafish

## Abstract

Cell ablation is a powerful method for elucidating the contributions of individual cell populations to embryonic development and tissue regeneration. Targeted cell loss in whole organisms has been typically achieved through expression of a cytotoxic or prodrug-activating gene product in the cell type of interest. This approach depends on the availability of tissue-specific promoters, and it does not allow further spatial selectivity within the promoter-defined region(s). To address this limitation, we have developed ablative methods that combine genetically encoded toxins, the tissue specificity afforded by *cis*-regulatory elements, and the conditionality of optogenetics. Using this integrative approach, we have ablated cells in zebrafish embryos with spatial and temporal precision.

## INTRODUCTION

Tissue development, homeostasis, and regeneration rely on functional networks composed of diverse cell types. To probe these mechanisms, several genetically encoded technologies have been developed to achieve cell ablation with spatiotemporal control, including cytotoxic proteins (Chelur and Chalfie, 2007; Kurita et al., 2003; Slanchev et al., 2005; Smith et al., 2002; Smith et al., 2007) and enzymes that can covert prodrugs into cytotoxic molecules (Bridgewater et al., 1995; Chu et al., 2007; Curado et al., 2007; Jung et al., 2007; Pisharath et al., 2007; Springer and Niculescu-Duvaz, 2000). Both of these approaches can be combined with tissue-specific promoters to spatially restrict ablation; however, *cis*-regulatory sequences that are exclusive to the targeted cells are not always available, resulting in collateral damage to other tissues. For example, nitroreductase and nitrofuran-based toxins have been used to ablate motor neurons in zebrafish larvae (Ohnmacht et al., 2016), providing an alternative to mechanically induced spinal cord lesions (Becker and Becker, 2001; Briona et al., 2015; Mokalled et al., 2016; Reimer et al., 2009; Yu et al., 2011). While this chemical-genetic technique was able to elicit regenerative responses in the central nervous system (CNS), the promoter used for these studies also led to nitroreductase expression in the heart and pancreas. As result, metronidazole treatment also induced heart damage and oedema (Ohnmacht et al., 2016).

Light-inducible technologies have the potential to achieve targeted cell ablation in a more precise manner. Toward that goal, chromophores that generate reactive oxygen species (ROS) upon exposure to light have been applied in various systems (Buckley et al., 2017; Bulina et al., 2006; Makhijani et al., 2017; Qi et al., 2012; Sarkisyan et al., 2015; Xu and Chisholm, 2016). However, these methods can require sustained irradiation for efficient cell ablation, limiting their ability to target specific cell populations in dynamic systems, and ROS sensitivity varies between cell types (Williams et al., 2013). We reasoned that light-inducible expression of cytotoxic proteins or toxin-activating enzymes might represent a more general and versatile approach. In particular, we envisioned this could be achieved using a photoactivatable transcription factor that can drive the expression of exogenous transgenes. One candidate construct for this strategy is a synthetic transcription factor, GAVPO, that was originally developed for light-inducible gene expression in mammalian systems (Wang et al., 2012). The GAVPO transactivator contains a Gal4 DNA-binding domain, a variant of the VIVID light-oxygen-voltage (LOV) domain, and the p65 transcriptional activation domain. Upon exposure to blue light, the LOV domain undergoes a photo-reductive reaction that forms a covalent adduct between its flavin adenine dinucleotide chromophore and a neighboring cysteine residue. The adduct stabilizes a conformational state that promotes LOV domain self-association, generating a transcriptionally active GAVPO dimer. Light-activated GAVPO then drives the expression of upstream activating sequence (UAS)-controlled genes until it reverts to the inactive monomer with a half-life of approximately 2 hours (Wang et al., 2012).

In this report, we investigate the efficacy of light-inducible cell ablation in zebrafish using two genetically encoded cell-ablation technologies and the GAVPO transactivator. We demonstrate that the cytotoxic ion channel variant M2^H37A^ (Lam et al., 2010; Smith et al., 2002; Smith et al., 2007) acts through non-apoptotic pathways to kill neurons and other cell types with greater efficacy and faster kinetics than the nitroreductase/nitrofuran system. We further establish GAVPO as an effective tool for achieving photoactivatable gene transcription in zebrafish, using both embryos injected with *GAVPO* mRNA and stable transgenic lines with neuronal expression of the transactivator. Finally, we show that integrating the GAVPO and M2^H37A^ technologies enables targeted cell ablations in zebrafish embryos and larvae. We anticipate that optogenetic cell ablation will be an invaluable experimental methodology for studying developmental and regenerative biology.

## RESULTS

### Genetically encoded neuron ablation in zebrafish

To develop conditional models of CNS damage, we pursued neuronal ablation through two genetically encoded technologies: bacterial nitroreductase (NTR) and viral ion channel M2. The *Escherichia coli* gene, *nsfB*, encodes a flavoprotein that can reduce a variety of nitroaromatic compounds (Zenno et al., 1996). Among the substrates of this nitroreductase (NTR) is the prodrug metronidazole, which is reductively converted into a DNA interstrand crosslinking agent that triggers cell death. M2 normally functions as a proton-selective channel during influenza infection (Shimbo et al., 1996; Wharton et al., 1994), and a single point mutation in the transmembrane domain (H37A) converts M2 into a constitutively-active, non-selective cation channel (Gandhi et al., 1999; Wang et al., 1995). This loss of specificity renders M2^H37A^ cytotoxic (Lam et al., 2010; Le Tissier et al., 2005; Smith et al., 2002; Smith et al., 2007). The availability of M2-targeting antiviral drugs, such as rimantadine and amantadine (Intharathep et al., 2008; Schnell and Chou, 2008), also enables pharmacological control of M2^H37A^ function (Fig. S1A).

Exogenous expression of NTR in combination with metronidazole has been widely used to selectively ablate cells in zebrafish (Curado et al., 2007; Curado et al., 2008; Pisharath and Parsons, 2009; Pisharath et al., 2007). However, the M2^H37A^ channel has not been applied previously in this model organism. We therefore injected zebrafish embryos with varying amounts of *M2*^*H37A*^ mRNA and monitored their embryonic development. Toxicity was initially observed at the blastula stages, and the embryos were unable to proceed through gastrulation (Fig. S1B-C). Injecting increasing amounts of *M2*^*H37A*^ mRNA rapidly killed 100% of the embryos. Culturing the M2^H37A^-expressing embryos in the presence of 100 μg/mL rimantadine enabled some to develop normally through 24 hours post fertilization (hpf). However, the channel blocker was unable to fully protect against channel-mediated toxicity, demonstrating the need to control *M2*^*H37A*^ expression.

We next examined whether M2^H37A^ can be used to ablate zebrafish neurons by establishing a system for Gal4/UAS-dependent expression in these cells. We generated heterozygous *Tg(UAS:M2^H37A^;myl7:mCherry)* zebrafish, using five tandem UAS sequences to minimize basal M2^H37A^ expression (Ma et al., 2013), and crossed them with a homozygous *Tg(tuba1a:Gal4VP16;myl7:GFP)* line. We then selected progeny that contained one copy of each transgene, which could be confirmed by the presence of both GFP and mCherry fluorescence in the heart (Fig. 1A). The double transgenic embryos expressed M2^H37A^ in neural tissues (Fig. S2), and they exhibited CNS deficits that appeared as early as 24 hpf and progressed over time (Fig. 1A). Rimantadine rescued these defects, confirming the role of M2^H37A^ channel activity in neuronal loss (Fig. 1A).

**Fig. 1.**
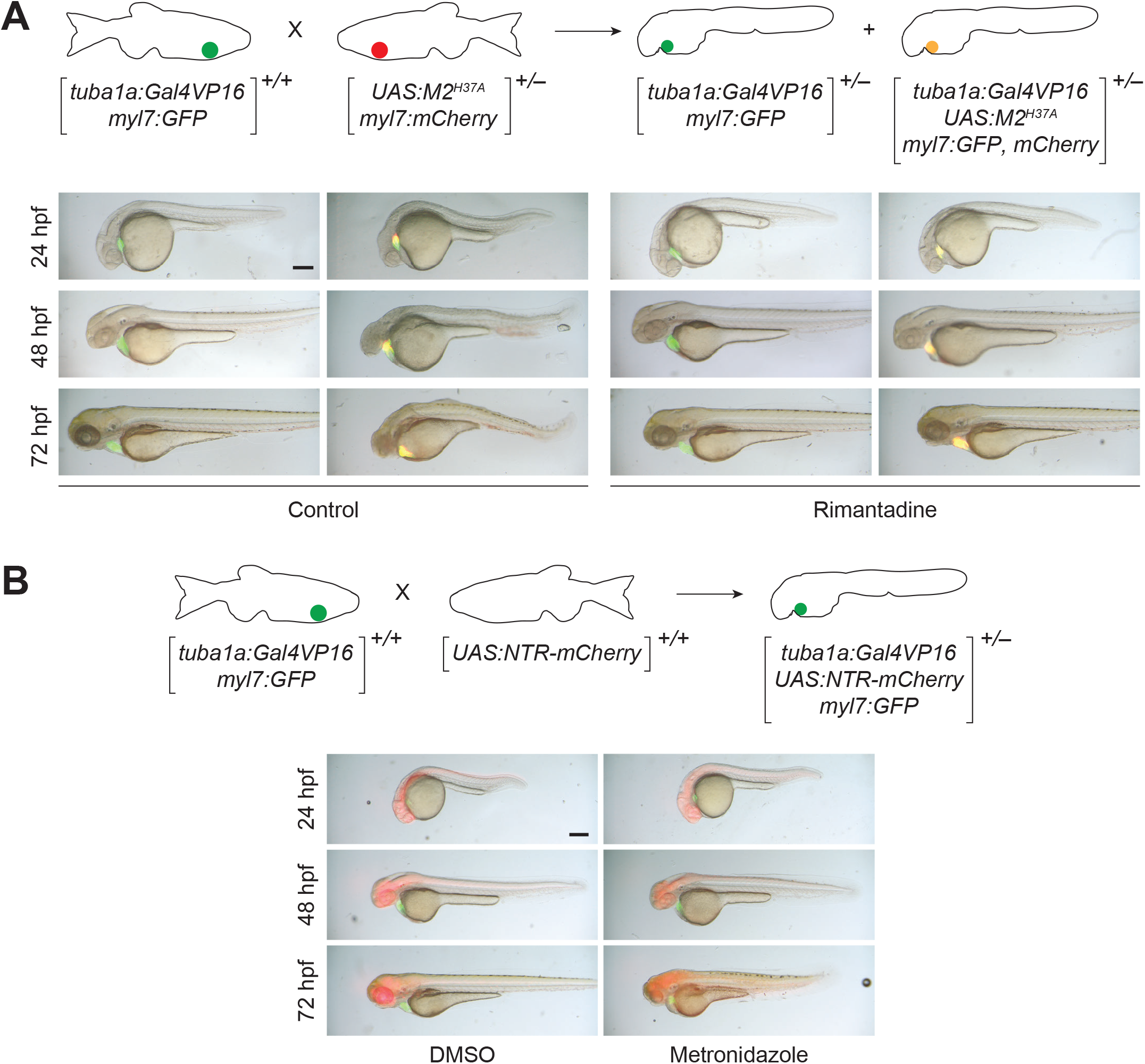
M2^H37A^ ablates neurons more rapidly than NTR. **(A)**. Comparison of *Tg(tuba1a:Gal4VP16;myl7:GFP)* and *Tg*(*tuba1a:Gal4VP16;UAS:M2^H37A^;myl7:gfp,mCherry*) embryos cultured in the absence or presence of 100 μg/mL rimantadine. Neuronal expression of M2^H37A^ induces loss of the midbrain-hindbrain boundary at 24 hpf and increasingly severe CNS deficits as development continues. Treatment with rimantadine starting at 10 hpf rescues these neuronal defects in *Tg*(*tuba1a:Gal4VP16;UAS:M2^H37A^;myl7:gfp,mCherry*) embryos. Representative brightfield and epifluorescence micrographs (overlays) from three independent experiments are shown. **(B)** Comparison of *Tg*(*tuba1a:Gal4VP16;UAS:NTR-mCherry;myl7:GFP*) embryos cultured with 5 mM metronidazole or an equivalent amount of DMSO, starting at 10 hpf. Metronidazole-treated embryos first exhibit CNS defects at 48 hpf. Representative brightfield and epifluorescence micrographs (overlays) from two independent experiments are shown. All embryos are shown in lateral view, anterior left. Scale bars: 250 μm. Statistics for the observed phenotypes are in Table S1.

To compare the ablative efficacies of the M2^H37A^ and NTR/metronidazole systems, we also crossed homozygous *Tg(UAS:NTR-mCherry)* (Goldman et al., 2001) and homozygous *Tg(tuba1a:Gal4VP16;myl7:GFP)* zebrafish and cultured the resulting embryos in medium supplemented with DMSO or 5 mM metronidazole starting at bud stage (10 hpf). In contrast to zebrafish with neuronal M2^H37A^ expression, metronidazole treatment of the NTR-expressing progeny did not induce CNS defects until 48 hpf, suggesting the two ablative technologies kill neural cells with different kinetics and possibly through distinct mechanisms (Fig. 1B). We next used TUNEL staining to detect apoptosis in M2^H37A^- and NTR-expressing embryos (Fig. S3A). Although we observed overt developmental defects as early as 24 hpf in M2^H37A^-expressing embryos, significant TUNEL staining was not detected until 32 hpf, and TUNEL-positive cells were distributed in regions within and near the CNS. NTR-expressing embryos treated with 5 mM metronidazole exhibited robust TUNEL staining localized to the CNS at 48 hpf, consistent with observed phenotypic defects. Thus, M2^H37A^ appears to kill zebrafish neurons through necrosis, which subsequently causes apoptosis of surrounding cells. This progressive mechanism of tissue loss is similar to that associated with CNS damage (Oyinbo, 2011), making M2^H37A^ a useful tool for studying neural injury and recovery.

### Light-inducible cell ablation using transient GAVPO expression

Using cytotoxic proteins to ablate specific cell populations, including regions within a tissue of interest, requires spatiotemporal control of their expression or function. To accomplish this goal, we pursued light-dependent NTR and M2^H37A^ expression in zebrafish embryos and larvae. A number of strategies for photoactivatable gene expression have been described, including GAVPO, EL222, an EL222-derived construct (TAEL), and two-hybrid-like systems based on CYR2-CIB1 or PhyB-PIF3 heterodimerization (Liu et al., 2012; Motta-Mena et al., 2014; Reade et al., 2017; Shimizu-Sato et al., 2002). Of these technologies, EL222, TAEL, and CYR2-CIB1have been applied previously in zebrafish, but they are limited by the minutes-scale lifetimes of their light-activated states. The PhyB-PIF3 system requires co-administration of phycocyanobilin, an exogenous chromophore that cannot penetrate zebrafish embryos (Buckley et al., 2016). Since GAVPO utilizes endogenous flavin and has a light-state half-life of 2 hours, we envisioned that this optogenetic tool could enable transcriptional control with transient illumination, a capability that would be particularly valuable for genetically encoded ablation of dynamic cell populations.

We examined GAVPO activity in *Tg(UAS:NTR-mCherry)* zebrafish by injecting transgenic zygotes with various amounts of *GAVPO* mRNA (Fig. 2B). The embryos were then raised in the dark until 6 hpf, after which half were exposed to blue light (470 nm) from a light-emitting diode (LED) lamp for 2 hours. NTR-mCherry levels were subsequently assessed by fluorescence microscopy at 24 hpf. GAVPO expression induced ubiquitous NTR-mCherry expression in a light- and dose-dependent manner (Fig. 2B), and the majority of animals developed normally when the *GAVPO* mRNA was injected at a dose of 100 pg/embryo (Fig. S4A). At 200 pg/embryo, *GAVPO* mRNA toxicity was similar to that observed with injection of 25 pg/embryo *Gal4VP16* mRNA (Fig. S4A). Taken together, these findings indicate that the GAVPO system can be used to convey light-dependent gene expression in multiple zebrafish cell types and may provide a less-toxic alternative to the currently available Gal4 driver lines.

**Fig. 2.**
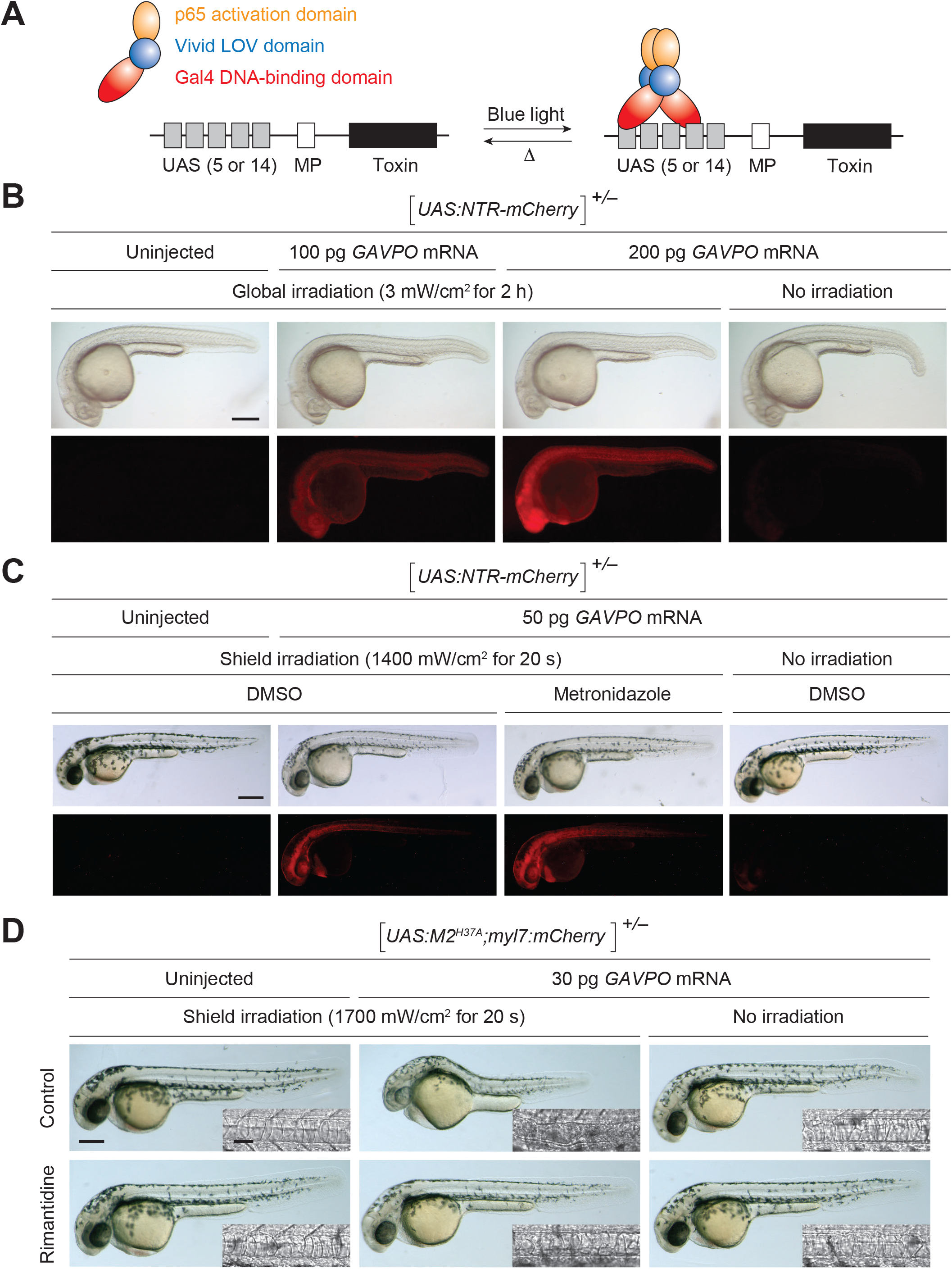
GAVPO conveys light-inducible gene expression in zebrafish. **(A)** Mechanism of GAVPO-dependent gene expression. **(B)** *Tg(UAS:NTR-mCherry)* zygotes were injected with the designated amounts of *GAVPO* mRNA, irradiated with a blue LED lamp at 6 hpf, and then imaged at 24 hpf. Representative brightfield and epifluorescence micrographs from two to four independent experiments are shown. GAVPO induces NTR-mCherry expression in a light- and concentration-dependent manner. **(C)** *Tg(UAS:NTR-mCherry)* zygotes were injected with the designated amounts of *GAVPO* mRNA, and their embryonic shields were irradiated at 6 hpf using an epifluorescence microscope equipped with a 470/40-nm filter. An iris diaphragm was used to restrict illumination to 100-μm-diameter region. The 10 hpf embryos were dechorionated and treated with DMSO or 5 mM metronidazole and imaged at 32 hpf. Representative brightfield and epifluorescence micrographs from two independent experiments are shown. Shield-irradiation of the GAVPO-expressing embryos induces NTR-mCherry expression in the notochord, head mesoderm, and hatching gland. **(D)** *Tg(UAS:M2^H37A^;myl7:mCherry)* zygotes were injected with the designated amounts of *GAVPO* mRNA, cultured in the absence or presence of 100 μg/mL rimantadine, shield-irradiated at 6 hpf as described in (C), and imaged at 30 hpf. Representative brightfield and differential interference contrast (DIC) micrographs from three independent experiments are shown. Shield-irradiation of the GAVPO-expressing embryos disrupts development of the notochord (inset), head, and hatching gland. All embryos are shown in lateral view, anterior left. Scale bars: B-D, 250 μm; D (inset), 50 μm. Statistics for the observed phenotypes are in Table S2.

We next examined whether GAVPO can be coupled with targeted illumination to achieve both spatial and temporal control of gene expression. We injected heterozygous *Tg(UAS:NTR-mCherry)* zygotes with *GAVPO* mRNA and irradiated the embryonic shield at 6 hpf for 20 seconds, using a epifluorescence microscope equipped with a 470/40-nm filter, 20x objective, and iris diaphragm. Zebrafish fate-mapping studies have shown that cells within this dorsal organizing center give rise to the notochord, head mesoderm, and hatching gland (Kimmel et al., 1990; Shih and Fraser, 1996), and we observed red fluorescence in these tissues at later developmental stages (Fig. 2C). To determine whether the expressed NTR-mCherry can impart nitrofuran sensitivity, we treated a cohort of the shield-irradiated embryos with either 5 mM metronidazole or DMSO vehicle alone from 10 to 32 hpf. We then fixed and immunostained the embryos for activated caspase-3, the initiating protease for the apoptotic pathway. Although basal levels of caspase-3 activation increased with metronidazole treatment, we did not observe any evidence of GAVPO/NTR-mediated apoptosis in the notochord (Fig. S3B). Thus, even levels of NTR-mCherry expression that can be readily detected by fluorescence microscopy are not sufficient to convey nitrofuran-induced apoptosis within this time frame.

Since our studies with *Tg(tuba1a:Gal4VP16;myl7:GFP)* zebrafish indicated that M2^H37A^ can ablate neurons with faster kinetics than the NTR/metronidazole system, we investigated whether the GAVPO transactivator could be used to express cytotoxic levels of this constitutively active channel. We injected varying amounts of *GAVPO* mRNA into heterozygous *Tg(UAS:M2^H37A^;myl7:mCherry)* zygotes, establishing 30 pg/embryo as the maximum dose that permits normal embryonic development in this transgenic line under dark-state conditions (Fig. S4B). Higher concentrations of *GAVPO* mRNA caused patterning defects, presumably due to the onset of light-independent M2^H37A^ expression. We then injected *Tg(UAS:M2^H37A^;myl7:mCherry)* zygotes with 30 pg/embryo of *GAVPO* mRNA and focally irradiated the shield with blue light at 6 hpf for 20 seconds using the epifluorescence microscope. The resulting embryos exhibited deformities of the notochord, head, and presumptive hatching gland when imaged at 30 hpf (Fig. 2D), matching the transgene expression we observed in *Tg(UAS:NTR-mCherry)* zebrafish that had been injected with *GAVPO* mRNA and shield-irradiated (Fig. 2C). In addition, culturing the focally irradiated embryos in the presence of the M2 blocker rimantadine, prevented these developmental defects.

### Light-inducible neuron ablation through tissue-specific GAVPO expression

Having established the ability of transiently expressed GAVPO to convey light-inducible cell ablation, we explored the efficacy of transgenic lines that stably express GAVPO in a neuron-specific manner. To target GAVPO expression to post-mitotic neurons, we generated zebrafish carrying *GAVPO* under control of the neuronal promoter *elavl3* (also known as *HuC*) (Kim et al., 1996; Park et al., 2000a; Park et al., 2000b). Multiple founders were screened for single insertions of the GAVPO expression construct and raised to generate two stable lines. When crossed to homozygous *Tg(UAS:NTR-mCherry)* zebrafish, both lines yielded embryos that exhibited photoinducible, neuron-specific mCherry fluorescence in the expected Mendelian ratios (Fig. S5A). One of the heterozygous lines exhibited more robust mCherry fluorescence in the developing spinal cord, and we incrossed this line to generate fertile, homozygous *Tg(elavl3:GAVPO)* zebrafish. *GAVPO* expression in this line recapitulated endogenous *elavl3* expression (Fig. S5B)

We next tested the ability of the *elavl3:GAVPO* transgene to convey light-inducible gene expression in neurons. We crossed homozygous *Tg(elavl3:GAVPO)* and homozygous *Tg(UAS:NTR-mCherry)* zebrafish to obtain embryos that contained one copy of each transgene. The double transgenic embryos were irradiated at different developmental stages for 2 hours using the blue LED lamp and imaged 8 hours post irradiation (hpi). We found that the mCherry fluorescence coincided with *GAVPO* expression at 36 and 48 hpf (Fig. S5B-C) and correlated with the duration and intensity of blue-light illumination (Fig. 3A-C). Focal irradiation using the epifluorescence microscope also induced mCherry expression in the targeted neurons within the zebrafish CNS (Fig. 3D-F).

**Fig. 3.**
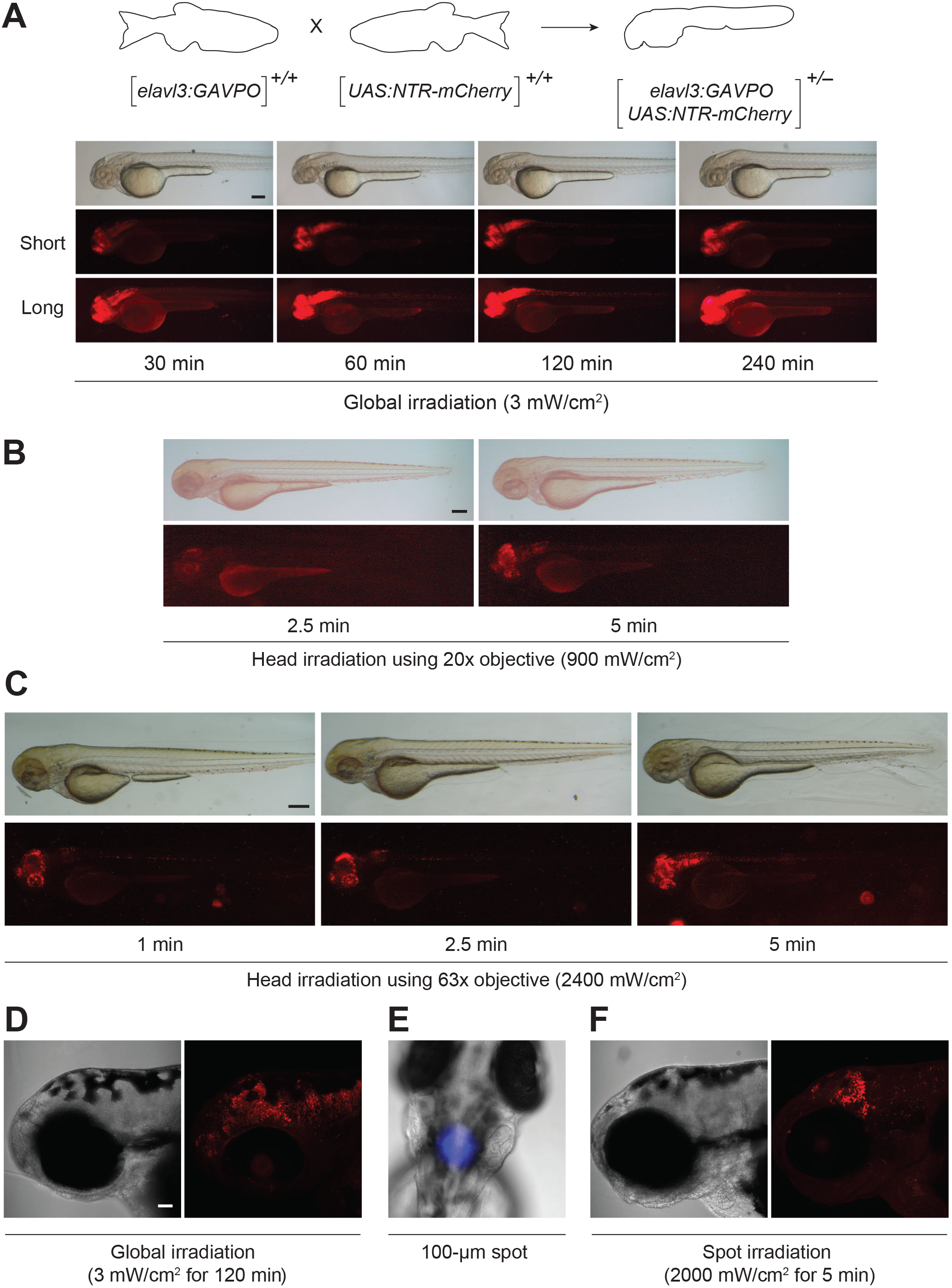
Light-inducible, neuron-specific gene expression using the GAVPO system. **(A)** *Tg(elavl3:GAVPO;UAS:NTR-mCherry)* embryos were irradiated for varying durations, using a blue LED lamp and starting at 48 hpf. The embryos were then imaged 8 hours later, and representative brightfield and epifluorescence micrographs (with short or long exposure times) are shown. NTR-mCherry was expressed in neural tissues, at levels that increased with the length of irradiation. **(B-C)** The heads of 48-hpf *Tg(elavl3:GAVPO;UAS:NTR-mCherry)* embryos were irradiated using an epifluorescence microscope equipped with a 470/40-nm filter and either a 20x (B) or 63x (C) objective. The embryos were imaged 8 hours later, and representative brightfield and epifluorescence micrographs are shown. NTR-mCherry expression was observed in anterior neural tissues, at levels that increased with the duration and intensity of irradiation. **(D-F)** *Tg(elavl3:GAVPO;UAS:NTR-mCherry)* embryos were illuminated either globally using a blue LED lamp (D) or within a 100-μm-diameter region in the head using epifluorescence microscope equipped with a 470/40-nm filter, 20x objective, and iris diaphragm (E-F), respectively. Representative brightfield and confocal fluorescence micrographs from two independent experiments are shown, and total sample sizes are indicated. LED illumination induced NTR-mCherry expression throughout the head, whereas spot-irradiated embryos exhibited mCherry fluorescence in only a localized region. Embryo orientations: A-D and F, lateral view and anterior left; E, dorsal view, anterior up. Scale bars: A, 200 μm; B-C, 300 μm; D-F, 50 μm. Statistics for the observed phenotypes are in Table S3.

We then crossed homozygous *Tg(UAS:M2^H37A^;myl7:mCherry)* zebrafish with the homozygous *Tg(elavl3:GAVPO)* line and exposed their *Tg(elavl3:GAVPO;UAS:M2^H37A^; myl7:mCherry)* progeny to blue light at 48 hpf for varying lengths of time. As determined by whole-mount immunostaining, M2^H37A^ levels correlated with the duration of blue-light irradiation (Fig. S6). Rimantadine treatment led to higher M2^H37A^ expression levels (Fig. S6), likely due to its protective effects on M2^H37A^-expressing neurons.

Finally, we sought to confirm that *Tg(elavl3:GAVPO;UAS:M2^H37A^;myl7:mCherry)* zebrafish exhibit light-dependent neuronal loss. Since M2^H37A^-induced cell death is not associated with caspase-3 activation or DNA fragmentation that can be detected by TUNEL staining, we monitored neuronal ablation by optical microscopy. We injected *Tg(elavl3:GAVPO;UAS:M2^H37A^; myl7:mCherry)* zygotes with an *elavl3:mCherry-CAAX* construct to label post-mitotic neurons in a mosaic manner (Fig. 4A), facilitating their visualization and quantification. We raised the embryos to 48 hpf, globally irradiated a subset with the blue LED lamp for 8 hours, and imaged the zebrafish at 12 hpi. The irradiated animals had significantly fewer intact neurons in the hindbrain than those maintained in the dark or cultured in rimantadine-containing medium after exposure to blue light (Fig. 4A). The spinal cords of irradiated *Tg(elavl3:GAVPO;UAS:M2^H37A^; myl7:mCherry)* embryos also exhibited reduced axon bundle widths (Fig. 4B).

**Fig. 4.**
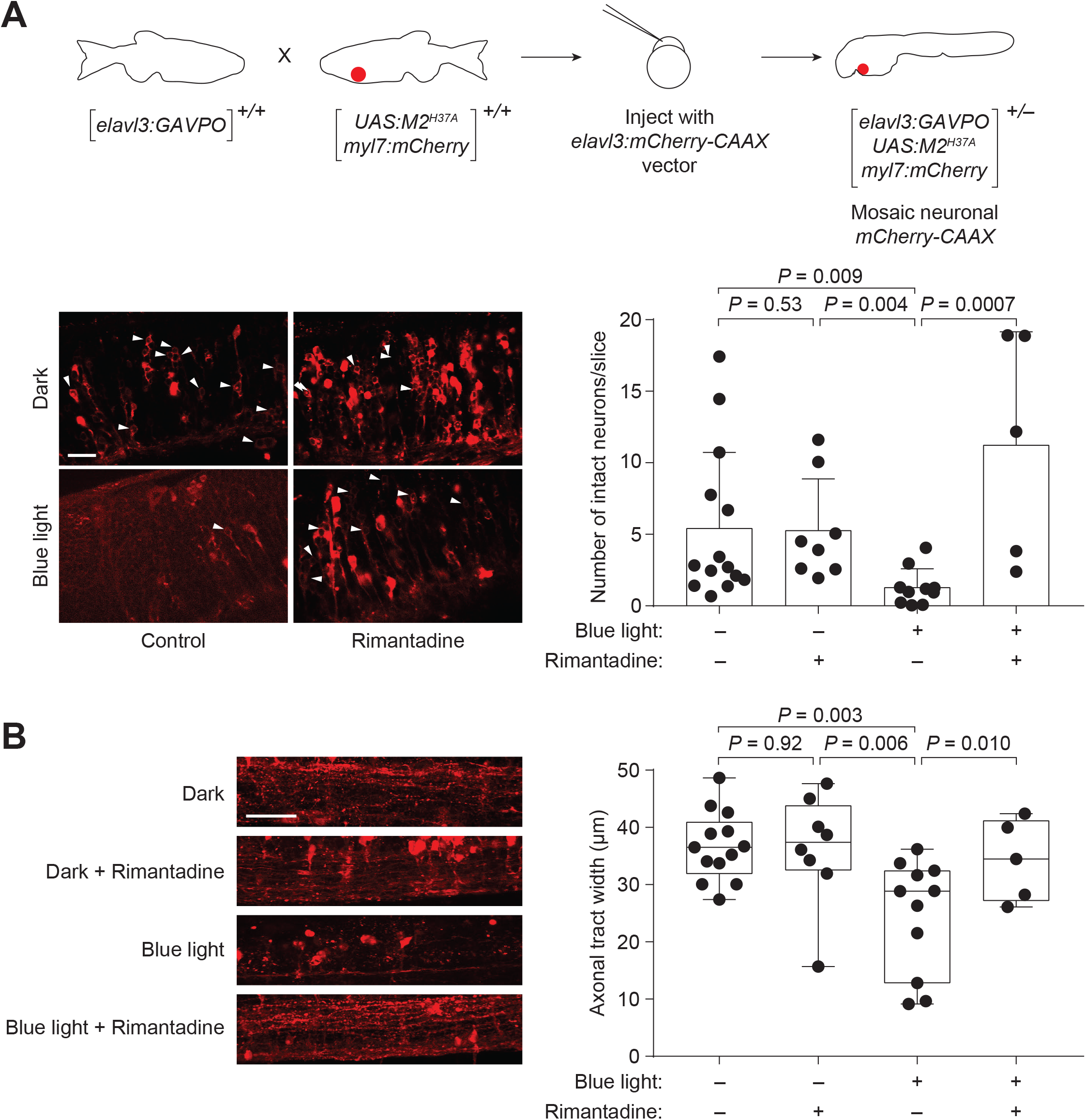
Light-inducible neural ablation using the GAVPO/M2^H37A^ system. **(A)** Heterozygous *Tg(elavl3:GAVPO; UAS:M2^H37A^;myl7:mCherry)* embryos were injected with an *elavl3:mCherry-CAAX* construct to label neurons in a mosaic manner. The injected embryos were irradiated with a blue LED lamp at 48 hpf for 8 hours, fixed 12 hours later, and then imaged on a confocal microscope. Representative micrographs from a hindbrain slice just above the otic vesicle are shown. Irradiated embryos have fewer intact neurons than those cultured in the dark or treated with 100 μg/mL rimantadine during blue-light illumination. Neurons were counted in ImageJ in a blinded manner, and the average number of neurons per slice ± s.d. for each experimental condition is shown. **(B)** Representative micrographs from a maximal intensity projection of the spinal cord just above the cloaca. The average axonal tract widths ± s.d. for each experimental condition are shown. Embryo orientations: A-B, Lateral view, anterior left. Scale bars: A-B, 25 μm. Values for individual fish are shown.

## DISCUSSION

Our studies demonstrate that GAVPO can be used to achieve optogenetic control of gene expression in zebrafish. Although it was previously reported that the *GAVPO* mRNA disrupts zebrafish development and reduces embryo viability (Reade et al., 2017), we readily generated viable, fertile transgenic lines, indicating that the photoactivatable transactivator is not inherently toxic. Accordingly, we observed that *GAVPO* mRNA is less toxic than *Gal4VP16* mRNA. The efficacy of GAVPO in zebrafish opens the door to new experimental approaches for this model organism, as this VIVID LOV domain-based system has certain advantages over other optogenetic approaches. For instance, the homodimeric GAVPO transactivator can be introduced into organisms as a single construct, whereas CYR2/CIB1- and PhyB/PIF-based technologies are heterodimeric systems (Buckley et al., 2016; Liu et al., 2012). The photoactivated state of GAVPO is also long-lived, with a mean lifetime that is approximately 360 times longer than that for EL222 and its derivatives (*τ* ~ 3 hours vs 30 seconds) (Motta-Mena et al., 2014; Wang et al., 2012)]. As a result, sustained GAVPO activity can be achieved with a single pulse of blue light, allowing transcriptional control of specific cell populations in developing embryos and other dynamic living systems. The reliance of GAVPO on endogenous flavin chromophores and its compatibility with the myriad of existing Gal4/UAS systems are other major strengths.

These attributes highlight the versatility of GAVPO as an optogenetic tool, and in principle, its functionality could be enhanced through modifications of its LOV domain. For example, the light-state lifetime of the VIVID LOV domain can be increased up to tenfold by mutating the flavin-binding site (Zoltowski et al., 2009). Introducing these residue changes into the GAVPO transactivator could prolong transcriptional responses to a single illumination pulse. We also observed that *GAVPO* mRNA was more toxic to *Tg(UAS:M2^H37A^;myl7:mCherry)* embryos than their *Tg(UAS:NTR-mCherry)* counterparts, even when the mRNA-injected zebrafish were cultured in the dark. Since M2^H37A^ is constitutively ablative and NTR is not, we interpret these results as evidence of dark-state GAVPO activity. Consistent with this model, rimantadine attenuates *GAVPO* mRNA-dependent toxicity in *Tg(UAS:M2^H37A^;myl7:mCherry)* embryos. We therefore anticipate that basal activation of UAS-driven transgenes will increase with GAVPO concentration, and strong promoters may constrain the dynamic range of this photoactivatable transactivator. This limitation could be overcome by using alternative *cis*-regulatory sequences, screening multiple transgenic lines, or alternatively, GAVPO variants with lower dark-state dimer affinities could be developed.

Our studies also establish M2^H37A^ expression as an effective strategy for ablating specific cell populations in whole organisms. In addition to providing the first demonstration that M2^H37A^ is cytotoxic in zebrafish, we reveal important differences between this non-selective cation channel and the NTR/metronidazole system. M2^H37A^ expression kills cells with greater efficacy than NTR-mediated prodrug activation, as evidenced by their respective activities in the zebrafish CNS and axial mesoderm. M2^H37A^ and NTR/metronidazole also ablate cells through distinct mechanisms. Neuronal M2^H37A^ expression appears to initially trigger necrosis, and surrounding cells can subsequently undergo apoptosis. In comparison, NTR-mediated activation of metronidazole induces programmed cell death. M2^H37A^-mediated cell ablation could therefore be a valuable method for modeling spinal cord injury, which involves necrosis and apoptosis (Oyinbo, 2011). Using M2^H37A^ to study molecular and cellular responses to CNS damage is particularly attractive, as commonly used models of spinal cord injury rely on mechanical perturbations that can concurrently damage muscle and skin (Becker et al., 1997; Bhatt et al., 2004). The sensitivity of M2^H37A^ to rimantadine also enables pharmacological tuning of this cytotoxic channel, allowing one to minimize basal activity and maximize inducibility.

Finally, by combining the GAVPO and M2^H37A^, we have achieved optical control of cell ablation in a whole organism. In comparison to other genetically encoded methods for cell ablation, the GAVPO/M2^H37A^ system can eliminate cells of interest without requiring tissue-specific promoters. When GAVPO is expressed ubiquitously, cell targeting can be guided by morphological cues alone, as illustrated by our ablation of progenitors within the zebrafish shield. The GAVPO/M2^H37A^ could be similarly applied to elucidate the developmental fates of other embryonic cell populations. Alternatively, *cis*-regulatory sequences can be used to restrict GAVPO expression to certain tissues such as the CNS, enabling the ablation of specific cell populations within these promoter-defined regions. The latter approach could be used to generate tissue injuries in a spatially precise manner. Based on these versatile capabilities, we anticipate that the GAVPO/M2^H37A^ system will be an invaluable tool for the deconstructing both developmental and regenerative mechanisms in whole organisms.

## MATERIALS AND METHODS

### Zebrafish husbandry

Adult zebrafish [wildtype AB strain and *Tg(UAS-E1b:NTR-mCherry)* (Davison et al., 2007); 3-18 months] were obtained from the Zebrafish International Resource Center. The transgenic line Tg(*tuba1a:Gal4VP16;myl7:GFP)* (Goldman et al., 2001) was a generous gift from Philippe Mourrain (Stanford University). All zebrafish lines were raised according to standard protocols. Embryos were obtained through natural matings and cultured at either 28.5 °C or 32 °C in E3 medium. The culture medium was also supplemented with 0.003% (w/v) PTU to prevent pigmentation. All animal procedures were approved by the Administrative Panel on Laboratory Animal Care (Protocol #10511) at Stanford University.

### Plasmids

To generate *GAVPO* mRNA, the *GAVPO* coding region with an upstream Kozak site was PCR-amplified from pGAVPO plasmid (gift from Yi Yang; East China University of Science and Technology) using the following primers containing BamHI and XbaI sites (underlined): 5-GAAT<u>GGATCC</u>GCCACCATGAAGCTACTGTC-3′ and 5′-GAAC<u>TCTAGA</u>GTGTACATTACTTGTCATCATCGTC-3′. The resulting PCR product was digested and ligated into pCS2+ to generate pCS2-GAVPO. The insertion was sequenced to confirm ligation fidelity. pCS2-M2^H37A^ (gift from Tim Mohun; Francis Crick Institute) and pCS2-GAVPO constructs were linearized with SacII, and their corresponding mRNAs were synthesized using *in vitro* SP6-dependent runoff transcription (Invitrogen).

To generate the Tol2-*elavl3:GAVPO* plasmid, the *GAVPO* coding region with an upstream Kozak sequence was PCR-amplified using the following primers: 5′-CCACCTGCAGATAATTGTTTAAACCACTCCGCCACCATGAAG-3′ and 5′-AGTAAAACG ACGGCCAGGATCCACCGGTCTGCTATTACTTGTCATCATCGTC-3′. Tol2-*elavl3:GCaMP6s* plasmid (Misha Ahrens; Addgene #59531) was digested with AgeI, and Tol2-*elavl3:GAVPO* was assembled from the resulting vector and GAVPO PCR product using Gibson Assembly Master Mix (NEB). The entire plasmid was sequenced to confirm ligation fidelity.

To generate the Tol2-*5xUAS-TATA:M2*^*H37A*^ Bleeding Heart construct, the *M2* coding region was PCR-amplified using the following primers which contain HindIII and ApaI sites (underlined): 5′-GAAC<u>AAGCTT</u>GCCACCATGAGTCTTCTAACCG-3′ and 5′-GAA<u>TGGGCCC</u>TTACTCCAGCTCTATGTTGAC-3′. This PCR product was then digested and ligated into pU5 (gift from Yi Yang; East China University of Science and Technology) to generate pU5-*M2^H37A^*. The *5xUAS-TATA:M2*^*H37A*^ region was then PCR-amplified using the following primers: 5′-ACACAGGCCAGATGTGGGCCCGTACTTGGAGCGGCCGCA-3′ and 5′-GTCTGGATCA TCATCGATGCGGCCGCAAGCCATAGAGCCCACCG-3′. pBH (gift from Michael Nonet; Washington University) was digested with ApaI and NotI, and the Tol2-*5xUAS-TATA:M2*^*H37A*^ Bleeding Heart plasmid was assembled from the digested pBH and *5xUAS-TATA:M2*^*H37A*^ PCR product using Gibson Assembly Master Mix (NEB). The entire plasmid was sequenced to confirm ligation fidelity.

To generate the Tol2-*elavl3:mCherry-CAAX* plasmid, the mCherry and *CAAX* coding regions with an upstream Kozak sequence was PCR amplified using the following primers. mCherry: 5′-TATATTTTCCACCTGCAGATAATTACCGGTGCCACCATGGTGAGCAAG-3′ and 5′-TAAAACGACGGCCAGGATCCACGCGTCTATTACATAATTACACACTTTGTCTTTGACTTC-3′; CAAX: 5′-TATATTTTCCACCTGCAGATAATTACCGGTTCCGCCACCATGGTGAGC-3′ and 5′-AGTAAAACGACGGCCAGGATCCACACGCGTCTATTACATAATTA CACACTTTGTCTTTGACTTCTTTTTCTTCTTTTTAC-3′. Tol2-*elavl3:GCaMP6s* was digested with AgeI, and Tol2-*elavl3:mCherry-CAAX* was assembled from the digested plasmid and PCR products using Gibson Assembly Master Mix (NEB). The insert was sequenced to confirm ligation fidelity.

### Generation of transgenic zebrafish lines

Transgenic zebrafish were generated using Tol2-mediated transgenesis as previously described (Suster et al., 2009). For each line generated, plasmid DNA and *Tol2* mRNA were premixed and co-injected into one-cell-stage embryos (12–50 pg of plasmid; 50 pg of mRNA). Fish were raised to adulthood and mated with wildtype AB fish to identify founders with germline transmission, and those yielding F2 generations with monoallelic expression were used to establish transgenic lines. Both heterozygous and homozygous *Tg*(*elavl3:GAVPO*) and homozygous *Tg*(*UAS:M2^H37A^;myl7:mCherry*) lines were used in subsequent studies.

### GAVPO photoactivation

All experimental procedures with GAVPO-expressing zebrafish utilized a Wratten #29 filter (Kodak) to minimize GAVPO activation by the microscopy light sources required for embryological procedures, monitoring, and imaging. For global irradiations, a plate containing E3 medium and embryos was mounted onto a mirrored surface and then fixed onto a Vortex-Genie (Scientific Industries). Dechorionated embryos were irradiated using a blue LED light source (TaoTronics). Light intensity received by the embryos was measured as 3 mW/cm^2^. For individual irradiations, the embryos were either mounted in an injection tray or in 1% low-melt agarose on a slide. Each embryo was irradiated using a Leica DM4500B upright compound microscope equipped with a mercury lamp, a GFP filter (ex: 470 nm, 40-nm bandpass), and a 20x/0.5 NA or a 63x water-immersion objective. For focal irradiation, an iris diaphragm was used to limit the targeted region to a 100-μm diameter. Light intensities from the mercury lamp were measured before and after the experiment, and they ranged from 900 to 2400 mW/cm^2^.

### Immunostaining

Embryos were fixed at the desired timepoint in 4% (w/v) paraformaldehyde (PFA) in PBS at 4 °C overnight and then stored in methanol. After rehydration, embryos were permeabilized with acetone, immunostained, and imaged with a Leica DM4500B compound microscope equipped with a Retiga-SRV or Prime cMOS camera (QImaging) or Zeiss LSM 700 and LSM 800 confocal microscopes equipped with a MA-PMT. The following antibodies were used: mouse monoclonal anti-HuC/HuD (1:100 dilution, Molecular Probes A-21271), mouse monoclonal anti-mCherry (1:500 dilution, Abcam ab125096), rabbit polyclonal anti-active Caspase-3 (1:100 dilution, BD Biosciences BDB559565), rabbit polyclonal anti-influenza A virus M2 (1:100 dilution, GeneTex GTX125951), goat anti-mouse and goat anti-rabbit antibodies conjugated to Alexa Fluor 488, 555, or 594 (1:200 dilution, Roche).

### TUNEL staining

For M2^H37A^ and NTR experiments, embryos were raised in E3 medium supplemented with either 100 μg/mL rimantadine or 5 mM metronidazole, respectively. Embryos raised in E3 medium treated with an equivalent amount of DMSO vehicle were used as controls. The embryos were then dechorionated using forceps, fixed in 4% PFA at the desired timepoint, and stored in methanol. Embryos were rehydrated and permeabilized with diluted bleach solution containing 0.8% potassium hydroxide, 3% hydrogen peroxide and 0.2% Triton X-100. After bleaching, the embryos were washed with PBS supplemented with 0.2% Triton X-100 (PBS-X), dehydrated with methanol, and stored overnight at –20 °C. After rehydration, embryos were repeatedly washed with PBS-X and digested using proteinase K. Embryos were re-fixed in 4% PFA followed by treatment with a pre-chilled solution (2:1) of ethanol and acetic acid. Using the ApopTag Peroxidase In Situ Apoptosis Detection Kit (Millipore Sigma), the samples were incubated in equilibration buffer for 1 hour and then TdT reaction buffer overnight at 37 ºC. Embryos were washed in diluted stop solution and PBS-X and then incubated with a blocking solution containing: 2% (w/v) blocking reagent (Roche), 0.15 M NaCl, 0.1 M maleic acid, and 20% (v/v) sheep serum (Sigma #S2263). The embryos were incubated with horseradish peroxidase-conjugated anti-digoxigenin antibody overnight at 4 °C and washed with PBS-X. Embryos were stained with fresh DAB mix (Metal Enhanced DAB Kit Thermo Fisher Scientific #34065), washed and fixed in 4% PFA. The embryos were stored and imaged in a solution of 90% glycerol in PBS.

### *In situ* hybridization

Whole-mount *in situ* hybridization was performed according to standard protocols (Broadbent and Read, 1999). Zebrafish cDNA was prepared from RNA extracted from embryos as previously described (Payumo et al., 2015). T3-promoter containing PCR products were then amplified with the designated primers (T3 sequence underlined): *elavl3*: 5′-CACCTCACGCATCCTGGTAA-3′ and 5′-<u>ATTAACCCTCACTAAAG</u>GGATGGTCTTGAACGAGACCTGC-3′; *GAVPO* Probe 1: 5′-AAGAAAAACCGAAGTGCGCC-3′ and 5′-<u>ATTAACCCTCACTAAAG</u>GGATGTTTCATCTCGCACCGGAA-3′; *GAVPO* Probe 2: 5′-GCTCTGATTCTGTGCGACCT-3′ and 5′-<u>ATTAACCCTCACTAAAG</u>GGAATCAGCATGGGCTCAGTTGT-3′. RNA probes were *in vitro* transcribed from the PCR products using the MEGAscript T3 Transcription Kit (Ambion), substituting nucleotides from the kit with digoxigenin-UTP (Roche). For GAVPO staining, both probes were used simultaneously in separate tubes. Embryos were imaged using a Leica M205FA microscope equipped with a SPOT Flex color camera and images were captured using SPOT software (Molecular Devices).

### Neuron labeling and counting

Heterozygous *Tg(elavl3:GAVPO;UAS:M2^H37A^;myl7:mCherry)* zygotes were co-injected with plasmid DNA encoding *elavl3:mCherry-CAAX* (25 pg/embryo) and *Tol2* mRNA (50 pg/embryo). The resulting embryos were raised until 48 hpf, irradiated for 8 hours, and then fixed 12 hours after irradiation in 4% PFA for 1 hour at room temperature. The embryos were then mounted laterally in 1% low-melting agarose and imaged with a Zeiss LSM 700 confocal microscope equipped with a 63x/0.5 NA water-immersion objective. Z-stacks were generated from images taken at 2- to 5-micron intervals, using the following settings: 1024×1024 pixels, 8 speed, 4 averaging. Neurons were scored as intact when mCherry fluorescence was localized only to the plasma membrane and excluded from the cytoplasm. Counting was performed by an observer blinded to the experimental conditions. For spinal cord measurements, maximum intensity projections were made in ImageJ, and three measurements were made for each projection and averaged to determine the axonal tract width.

### Statistical analyses

For all zebrafish experiments, at least two breeding tanks, each containing 3 to 4 males and 3 to 5 females from separate stocks, were set up to generate embryos. Embryos from each tank were randomly distributed across tested conditions, and unfertilized and developmentally abnormal embryos were removed prior to irradiation or compound treatment. No statistical methods were used to determine sample size per condition. Experimental statistics for the phenotypes reported in Figures 1–3 are provided in Tables S1-S3, respectively. χ-square analyses were conducted for the phenotypic distributions reported in Figures S1 and S4. For the cell counting and axonal tract width measurements reported in Figure 4, the scorer was blinded to treatment conditions. Values for individual fish are plotted, and each distribution was assessed using the Shapiro-Wilk test and determined to be non-normal. A Kruskal-Wallis test with a two-stage linear step-up procedure of Benjamini, Krieger and Yekutieli was used to determine differences between all conditions. *P* values were corrected for multiple comparisons testing by controlling the false discovery rate to 0.05.

## ACKNOWLEDGMENTS

We gratefully acknowledge financial support from NIH R35 GM127030 (J.K.C.), a Craig H. Neilsen Postdoctoral Fellowship (K.M.), a Stanford School of Medicine Dean’s Postdoctoral Fellowship (P.C.), and the Stanford ChEM-H Undergraduate Scholars Program (P.A.P.). Imaging was supported in part by the University of Wyoming’s Integrated Microscopy Core (NIH P20 GM121310).

## AUTHOR CONTRIBUTIONS

K.M., P.C., and J.K.C. designed the experiments. K.M., P.C., P.A.P, and M.A.A. performed the experiments and analyzed data. K.M., P.C., and J.K.C. wrote the manuscript.

## SUPPORTING INFORMATION

Tables S1-S3 and Figs. S1-S6.

